# Evolution of Endothelin signaling and diversification of adult pigment pattern in *Danio* fishes

**DOI:** 10.1101/363879

**Authors:** Jessica E. Spiewak, Emily J. Bain, Jin Liu, Kellie Kou, Samantha L. Sturiale, Larissa B. Patterson, Parham Diba, Judith S. Eisen, Ingo Braasch, Julia Ganz, David M. Parichy

## Abstract

Fishes of the genus *Danio* exhibit diverse pigment patterns that serve as useful models for understanding the genes and cell behaviors underlying the evolution of adult form. Among these species, zebrafish *D. rerio* exhibit several dark stripes of melanophores with sparse iridophores that alternate with light interstripes of dense iridophores and xanthophores. By contrast, the closely related species *D. nigrofasciatus* has an attenuated pattern with fewer melanophores, stripes and interstripes. Here we demonstrate species differences in iridophore development that presage the fully formed patterns. Using genetic and transgenic approaches we identify the secreted peptide Endothelin-3 (Edn3)—a known melanogenic factor of tetrapods—as contributing to reduced iridophore proliferation and fewer stripes and interstripes in *D. nigrofasciatus*. We further show the locus encoding this factor is expressed at lower levels in *D. nigrofasciatus* owing to *cis*-regulatory differences between species. Finally, we show that functions of two paralogous loci encoding Edn3 have been partitioned between skin and non-skin iridophores. Our findings reveal genetic and cellular mechanisms contributing to pattern differences between these species and suggest a model for evolutionary changes in Edn3 requirements across vertebrates.

**Author Summary:** Neural crest derived pigment cells generate the spectacular variation in skin pigment patterns among vertebrates. Mammals and birds have just a single skin pigment cell, the melanocyte, whereas ectothermic vertebrates have several pigment cells including melanophores, iridophores and xanthophores, that together organize into a diverse array of patterns. In the teleost zebrafish, *Danio rerio*, an adult pattern of stripes depends on interactions between pigment cell classes and between pigment cells and their tissue environment. The close relative, *D. nigrofasciatus* has fewer stripes and prior analyses suggested a difference between these species that lies extrinsic to the pigment cells themselves. A candidate for mediating this difference is Endothelin-3 (Edn3), essential for melanocyte development in warm-blooded animals, and required by all three classes of pigment cells in an amphibian. We show that Edn3 specifically promotes iridophore development in *Danio*, and that differences in Edn3 expression contribute to differences in iridophore complements, and striping, between *D. rerio* and *D. nigrofasciatus*. Our study reveals a novel function for Edn3 and provides new insights into how changes in gene expression yield morphogenetic outcomes to effect diversification of adult form.

## Introduction

Mechanisms underlying species differences in adult form remain poorly understood. Quantitative genetic analyses and association studies have made progress in identifying loci, and even specific nucleotides, that contribute to morphological differences between closely related species and strains. Yet it remains often mysterious how allelic effects are translated into specific cellular outcomes of differentiation and morphogenesis to influence phenotype. Elucidating not only the genes but also the cellular behaviors underlying adult morphology and its diversification remains an outstanding challenge at the interface of evolutionary genetics and developmental biology.

To address genes and cellular outcomes in an evolutionary context requires a system amenable to modern methods of developmental genetic analysis and rich in phenotypic variation. Ideally the trait of interest would have behavioral or ecological implications, and its phenotype would be observable at a cellular level during development. In this context, adult pigment patterns of fishes in the genus *Danio* provide a valuable opportunity to interrogate genetic differences and the phenotypic consequences of these differences.

*Danio* fishes exhibit adult pigment patterns that include horizontal stripes, vertical bars, dark spots, light spots, uniform patterns and irregularly mottled patterns [1]. Pattern variation affects shoaling and might plausibly impact mate recognition, mate choice, and susceptibility to predation [2–5]. Phylogenetic relationships among species and subspecies are increasingly well understood, as is their biogeography, and some progress has been made towards elucidating their natural history [1,6–9]. Importantly in a developmental genetic context, one of these species, zebrafish *D. rerio*, is a well-established biomedical model organism with the genetic, genomic and cell biological tools that accompany this status. Such tools can be deployed in other danios to understand phenotypic diversification.

Adult pigment pattern formation in *D. rerio* is becoming well described in part because cellular behaviors can be observed directly in both wild-type and genetically manipulated backgrounds. The adult pigment pattern comprises three major classes of pigment cells—black melanophores, iridescent iridophores and yellow–orange xanthophores—all of which are derived directly or indirectly from embryonic neural crest cells [10,11]. The fully formed pattern consists of dark stripes of melanophores and sparse iridophores that alternate with light “interstripes” of xanthophores and dense iridophores (Figure 1, top). During a larva-to-adult transformation, precursors to adult iridophores and melanophores migrate to the skin from locations in the peripheral nervous system [10,12,13]. Once they reach the skin hypodermis, between the epidermis and the underlying myotome, the cells differentiate. Iridophores arrive first and establish a “primary” interstripe near the horizontal myoseptum [14–16]. Differentiating melanophores then form primary stripes dorsal and ventral to the interstripe, with their positions determined in part by interactions with iridophores. Later, xanthophores differentiate within the interstripe and these cells, as well as undifferentiated xanthophores, interact with melanophores to fully consolidate the stripe pattern [11,17–22]. As the fish grows, the pattern is reiterated: loosely arranged iridophores appear within stripes and expand into “secondary” interstripes where they increase in number and establish boundaries for the next forming secondary stripe [13,23]. Stripe development in *D. rerio* thus depends on serially repeated interactions among pigment cell classes. It also depends on factors in the tissue environment that are essential to regulating when and where pigment cells of each class appear [11,16].

**Figure 1.**
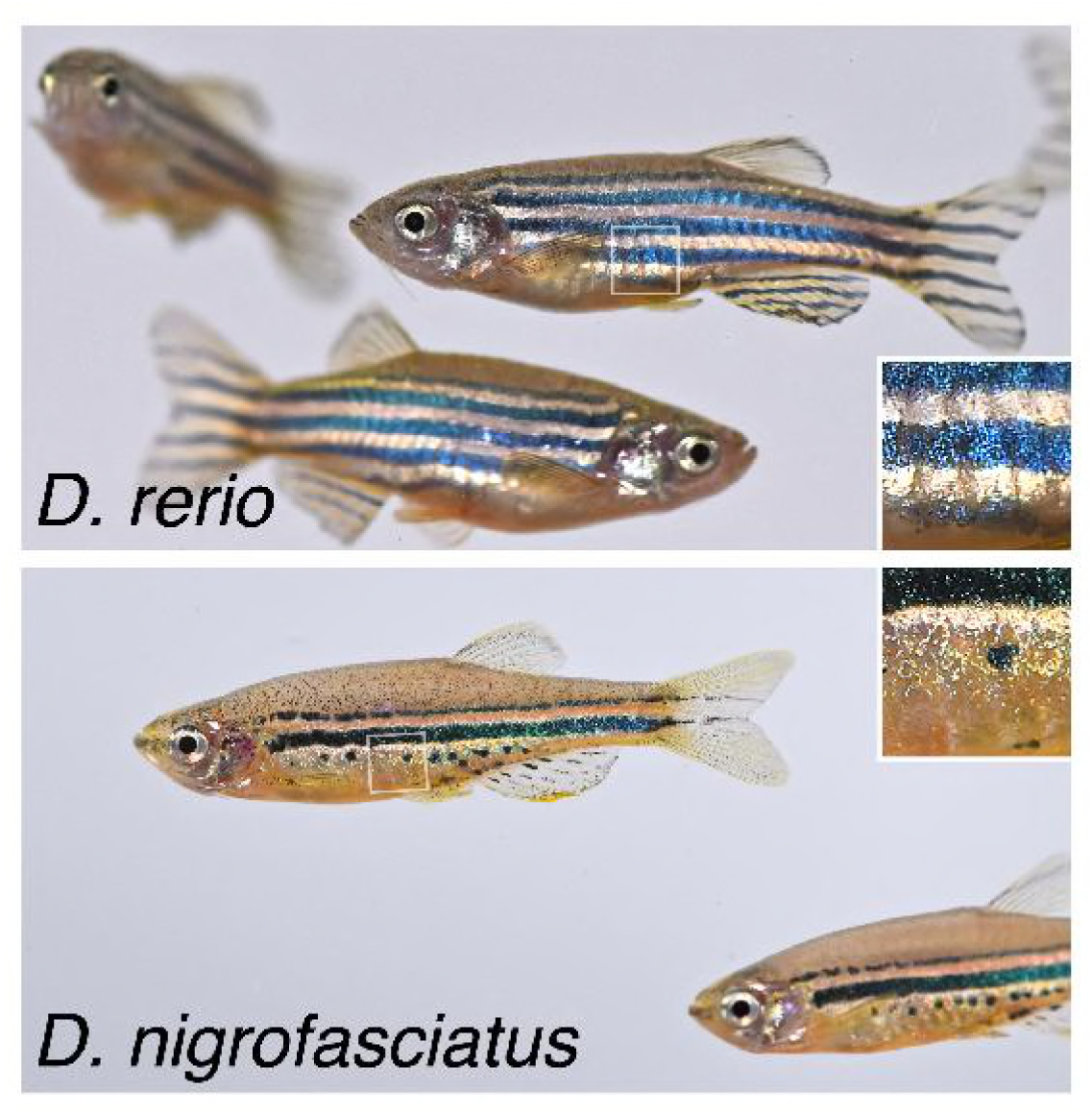
Different pigment patterns of *D. rerio* and *D. nigrofasciatus*. *Danio rerio* exhibit several dark stripes of melanophores with sparse iridophores, and light interstripes with abundant iridophores. *Danio nigrofasciatus* share common pattern elements but have fewer stripes and interstripes overall with spots forming ventrally instead of stripes. A shiny ventrum in both species results principally from iridophores that line the peritoneum, rather than iridophores in the hypodermis of the skin. Insets show iridescence of hypodermal iridophores.

Analyses of pattern development in other *Danio* are beginning to illuminate how pigment-cell “intrinsic” and “extrinsic” factors have influenced pattern evolution and the genetic bases for such differences [11,21,23–25]. Here, we extend these studies by examining pattern formation in *D. nigrofasciatus* (Figure 1, bottom). *D. rerio* and *D. nigrofasciatus* are closely related and occur within the “*D. rerio* species group” [9]. The essential elements of their patterns—stripes and interstripes—and the cell types comprising these patterns are the same. Nevertheless, *D. nigrofasciatus* has a smaller complement of adult melanophores than *D. rerio* and its stripes are fewer in number, with only residual spots where a secondary ventral stripe would form in *D. rerio*. Given the broader distribution of patterns and melanophores complements across *Danio*, the *D. nigrofasciatus* pattern of attenuated stripes is likely derived relative to that of *D. rerio* and other danios [26,27]. Cell transplantation analyses revealed that species differences in pattern result at least in part from evolutionary alterations residing in the extracellular environment that melanophores experience, rather than factors autonomous to the melanophores themselves.

In this study, we show that *D. rerio* and *D. nigrofasciatus* differ not only in melanophore complements but also iridophore behaviors. We show that iridophore development is curtailed in *D. nigrofasciatus*, with a corresponding loss of pattern reiteration. Using genetic and transgenic manipulations, we identify the endothelin pathway, and specifically the skin-secreted factor, Endothelin-3 (Edn3), as a candidate for mediating a species difference in iridophore proliferation. We find that *Danio* has two Edn3-encoding loci, arisen from an ancient genome duplication in the ancestor of teleost fishes [28,29], that have diverged in function to promote the development of different iridophore subclasses. One of these, *edn3b*, is required by hypodermal iridophores and has undergone *cis*-regulatory alteration resulting in diminished Edn3 expression in *D. nigrofasciatus*. Endothelin signaling is required directly by melanocytes in birds and mammals [30–33] but our findings indicate a specific role for Edn3b in promoting iridophore development, with only indirect effects on melanophores. These results suggest a model for the evolution of Edn3 function across vertebrates and implicate changes at a specific locus, *edn3b*, in altering cellular behavior that determines the numbers of stripes comprising adult pattern.

## Results

### Different iridophore complements of *D. nigrofasciatus* and *D. rerio*

Iridophores are essential to stripe reiteration of *D. rerio* [23] and iridophore-deficient mutants have fewer melanophores [15,16,34]. Given the fewer stripes and melanophores of *D. nigrofasciatus* (Figure 2A) [27], we asked whether iridophore development differs in this species from *D. rerio*. Figure 2B (upper) illustrates ventral pattern development of *D. rerio*. Iridophores were confined initially to the primary interstripe but subsequently occurred as dispersed cells further ventrally [13,16,23]. Additional melanophores developed ventrally to form the ventral primary stripe. Dispersed iridophores were found amongst these melanophores and, subsequently, additional iridophores developed further ventrally as the ventral secondary interstripe. In *D. nigrofasciatus*, however, very few dispersed iridophores developed ventral to the primary interstripe (Figure 2B, lower). Melanophores of the prospective ventral primary stripe initially occurred further ventrally than in *D. rerio* (also see [27]), similar to mutants of *D. rerio* having iridophore defects [16]. Few iridophores were evident either within the prospective ventral primary stripe or further ventrally.

**Figure 2.**
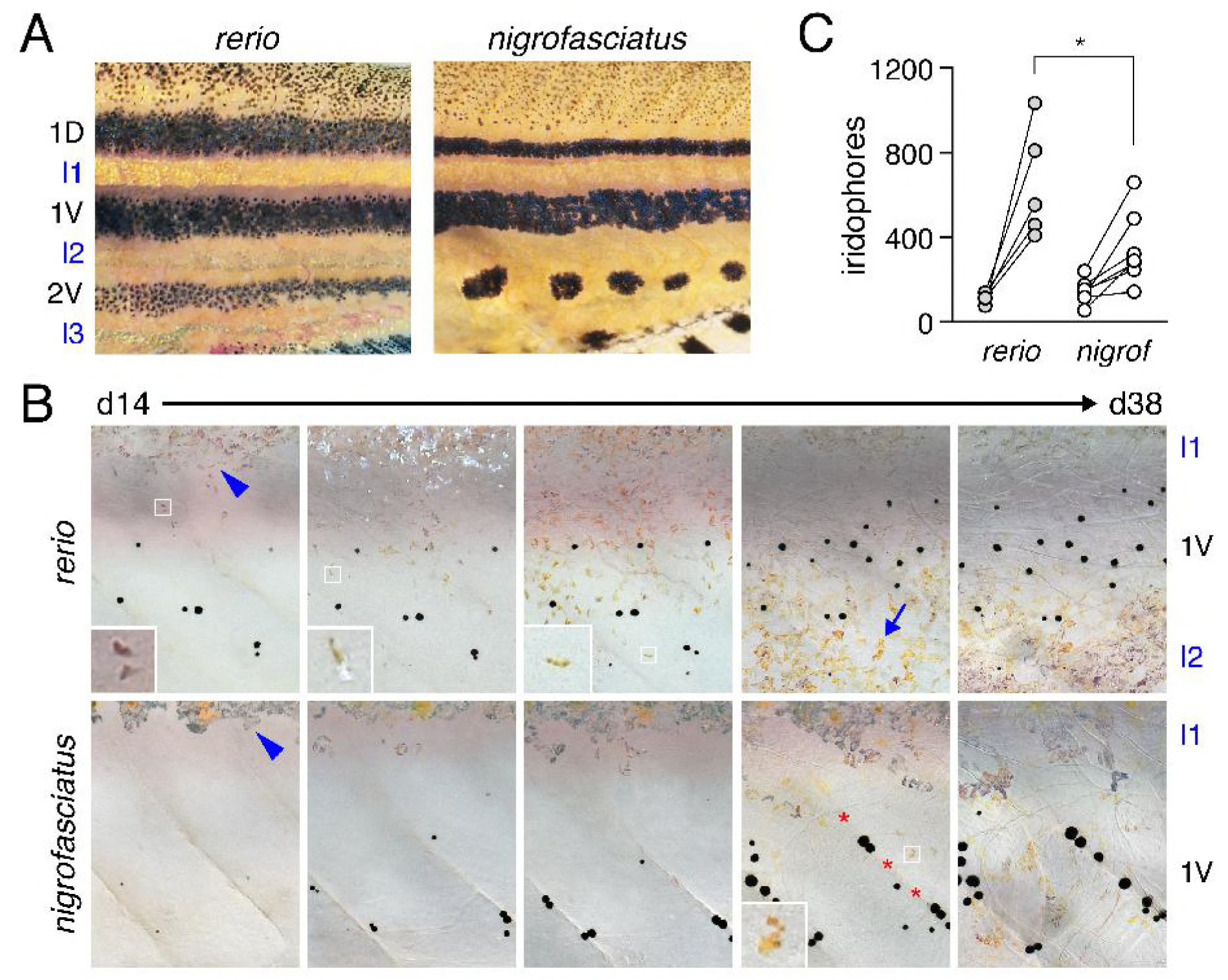
Iridophore development differs between *D. rerio* and *D. nigrofasciatus*. (A) Young adult patterns of the two species, illustrating fewer melanophores of *D. nigrofasciatus* compared to *D. rerio*. Stripes and interstripes are marked at the left. 1D, 1V: primary dorsal and ventral stripes. 2V, secondary ventral stripe. I1, I2, I3: Primary, secondary and tertiary interstripes. (B) Iridophores during primary stripe and secondary interstripe formation. Shown are representative individuals imaged repeatedly for *D. rerio* (upper) and *D. nigrofasciatus* (lower), with iridophores of the primary interstripe indicated by blue arrowheads. Fish were imaged throughout adult pattern formation with stages PB through J [14] illustrated here (corresponding to days ~14 through 38 post fertilization, shown for heuristic purposes only). Insets show iridophores at higher magnification. In *D. nigrofasciatus*, iridophores are comparatively few, do not as extensively populate the region of the secondary ventral stripe or the secondary ventral interstripe (blue arrow in *D. rerio*). Melanophores of the primary ventral stripe occur more ventrally than in *D. rerio* and tended to be more closely associated with vertical myosepta (marked by red asterisks). Sample sizes (*N*): 9 *D. rerio;* 6 *D. nigrofasciatus*. (C) Clonally related iridophores increased in number in both species between formation of primary interstripe (left; stage PB_+_) and subsequent pattern reiteration (right; J_++_). Points connected by lines represent individual at each developmental stage. Starting numbers were not significantly different (*F*_1,10_=0.94, *P*=0.4), whereas final numbers were significantly fewer in *D. nigrofasciatus* than in *D. rerio* (repeated measures, species x stage interaction, *F*_1,10_=7.47, *P*<0.05; *N*=5 *D. rerio*, *N*=6 *D. nigrofasciatus*).

Iridophores arise from progenitors that are established in association with the peripheral nervous system. These cells migrate to the hypodermis where they differentiate [12]. Individual progenitors can generate large hypodermal clones that expand during pattern formation [13]. To assess initial iridophore clone size and subsequent expansion we injected *D. rerio* and *D. nigrofasciatus* with limiting amounts of *pnp4a:palmEGFP* to drive membrane-targeted GFP in iridophores [21]. At transgene concentrations used, ~1% of injected embryos exhibited a single small patch of EGFP+ iridophores, consistent with labeling of individual progenitors [35,36]. Iridophore morphologies and initial clone sizes were similar between species, but subsequent expansion was significantly greater in *D. rerio* than *D. nigrofasciatus* (Figure 2C; Figure S1).

These observations indicate that adult pattern differences between *D. rerio* and *D. nigrofasciatus* are presaged not only by differences in melanophore development [27] but changes in iridophore behavior as well. This raises the possibility that evolutionary modifications to iridophore morphogenesis or differentiation have contributed to overall pattern differences between species.

### Endothelin pathway mutants identify a candidate gene for the reduced melanophore complement of *D. nigrofasciatus*

Shared phenotypes of laboratory induced mutants and other species identify candidate genes that may have contributed to morphological diversification [25,37,38]. *endothelin b1a receptor* (*ednrb1a*) mutant zebrafish resemble *D. nigrofasciatus* with deficiencies in iridophores and melanophores compared to wild-type *D. rerio*, and a pattern of stripes dorsally with spots ventrally. Prior genetic analyses failed to identify an obvious role for *ednrb1a* alleles in contributing to these species differences [37]. Ednrb1a is also expressed by pigment cells [34], whereas interspecific cell transplants suggested that pattern differences between *D. rerio* and *D. nigrofasciatus* likely result from differences in the tissue environment encountered by pigment cells [27]. Accordingly, we hypothesized that differences in expression of Ednrb1a ligand, Endothelin-3 (Edn3), contributes to the pigment pattern differences between these fishes. To first ascertain the phenotype of Edn3 mutants of *D. rerio* we induced mutations in each of two Edn3-encoding loci of zebrafish, *edn3a* (chromosome 11) and *edn3b* (chromosome 23) (Figure S2).

Fish homozygous mutant for an inactivating allele of *edn3a* exhibited relatively normal stripes and interstripes, but were deficient for iridophores that normally line the peritoneum, resulting in a rosy cast to the ventrum (Figure 3). By contrast, each of three *edn3b* presumptive null alleles exhibited severe deficiencies of hypodermal iridophores and melanophores and patterns of stripes breaking into spots; similar to *D. nigrofasciatus*, none had defects in peritoneal iridophores (Figure S3).

**Figure 3.**
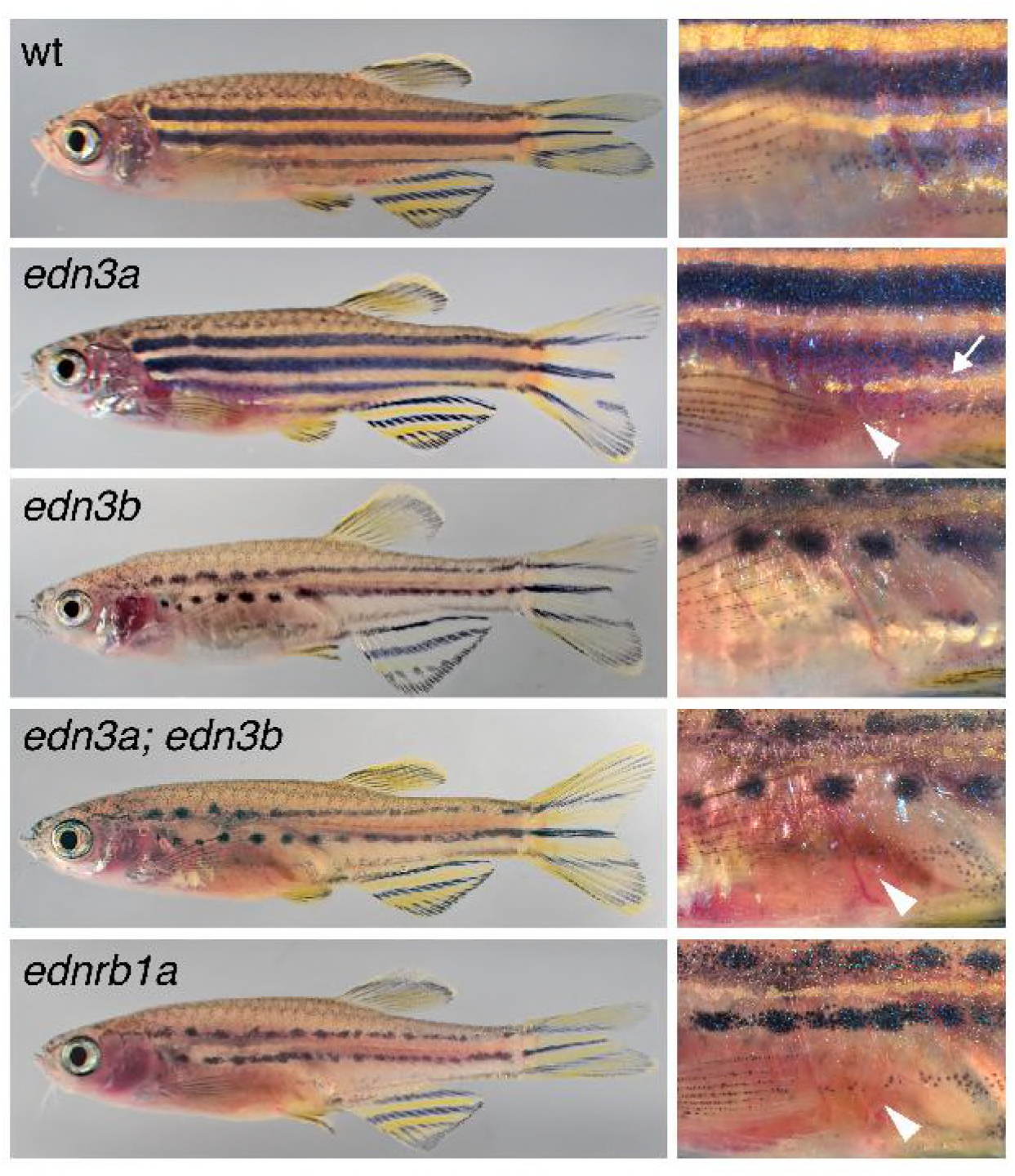
Edn3 and Ednrb1a mutants of *D. rerio*. Shown are wild-type (wt) and homozygous mutants for *edn3a* and *edn3b*, double mutant *edn3a; edn3b*, and *ednrb1a. edn3a* mutants had normal hypodermal pigment pattern, including iridophore interstripes (arrow, right) but lacked peritoneal iridophores (arrowhead). *edn3b* mutants had hypodermal iridophore and melanophore deficiencies but normal peritoneal iridophores. Fish doubly mutant for these loci exhibited both defects and resembled *ednrb1a* mutants.

*ednrb1a* mutants are defective for both hypodermal and peritoneal iridophores [34], suggesting that Edn3 signaling may have been partitioned evolutionarily between the two paralogous, ligand-encoding loci. Consistent with this idea, fish doubly mutant for *edn3a* and *edn3b* were deficient for both types of iridophores and resembled mutants for *ednrb1a* (Figure 3). These observations also suggest that Ednrb1a need only interact with Edn3a and Edn3b ligands to fulfill requirements for adult pigmentation, though Ednrb1 receptors of other vertebrate lineages are capable of transducing signals via other endothelins [29].

### Genetic analyses implicate *edn3b* in pattern difference between *D. rerio* and *D. nigrofasciatus*

The similarity of *edn3b* mutant *D. rerio* and *D. nigrofasciatus*—with fewer hypodermal melanophores and iridophores than wild-type *D. rerio*, but persisting peritoneal iridophores— identified *edn3b* as a particularly good candidate for contributing to the species difference in pigmentation. To assess this possibility further we used an interspecific complementation test [37–39]. If a loss-of-function *edn3b* allele contributes to the reduced iridophores and melanophores of *D. nigrofasciatus* compared to *D. rerio*, we would expect that in hybrids of *D. rerio* and *D. nigrofasciatus*, substitution of a *D. rerio* mutant *edn3b* (*edn3b*^*rerio*−^) allele for a *D. rerio* wild-type *edn3b* (*edn3b*^*rerio*+^) allele should expose the “weaker” *D. nigrofasciatus* allele, reducing the complement of iridophores and melanophores. Such an effect should be of greater magnitude than substituting a mutant for wild-type allele in *D. rerio*, and should be detectable as an allele x genetic background interaction. We therefore generated crosses of *edn3b*/+ *D. rerio* x *D. nigrofasciatus* as well as *edn3b*/+ x *edn3b*/+ *D. rerio*. We grew offspring until juvenile pigment patterns had formed, then genotyped individuals of hybrid (h) or *D. rerio* (r) backgrounds for the presence of either *edn3b*^*rerio*+^ or *edn3b*^*rerio*−^.

Hybrids between *D. rerio* and *D. nigrofasciatus* have patterns intermediate between the two species [37]. Figure 4A illustrates reduced coverage of iridophores and somewhat narrower stripes in fish carrying *edn3b*^*rerio*−^ as compared to siblings carrying *edn3b*^*rerio*+^. Total areas covered by interstripe iridophores were significantly reduced in hybrids compared to *D. rerio*, overall, and in both backgrounds by substitution of *edn3b*^*rerio*−^ for *edn3b*^*rerio*+^ (Figure 4B). Moreover, hybrids were more severely affected by this substitution than were *D. rerio*, resulting in a significant allele x genetic background interaction. Melanophore numbers were also reduced by substitution of *edn3b*^*rerio*−^ for *edn3b*^*rerio*+^ but hybrids were not significantly more affected than *D. rerio* (Figure 4C). These analyses suggest that the wild-type *D. nigrofasciatus edn3b* allele is hypomorphic to the wild-type *D. rerio* allele of *edn3b*, and support a model in which evolutionary changes at *edn3b* have affected iridophore coverage between species.

**Figure 4.**
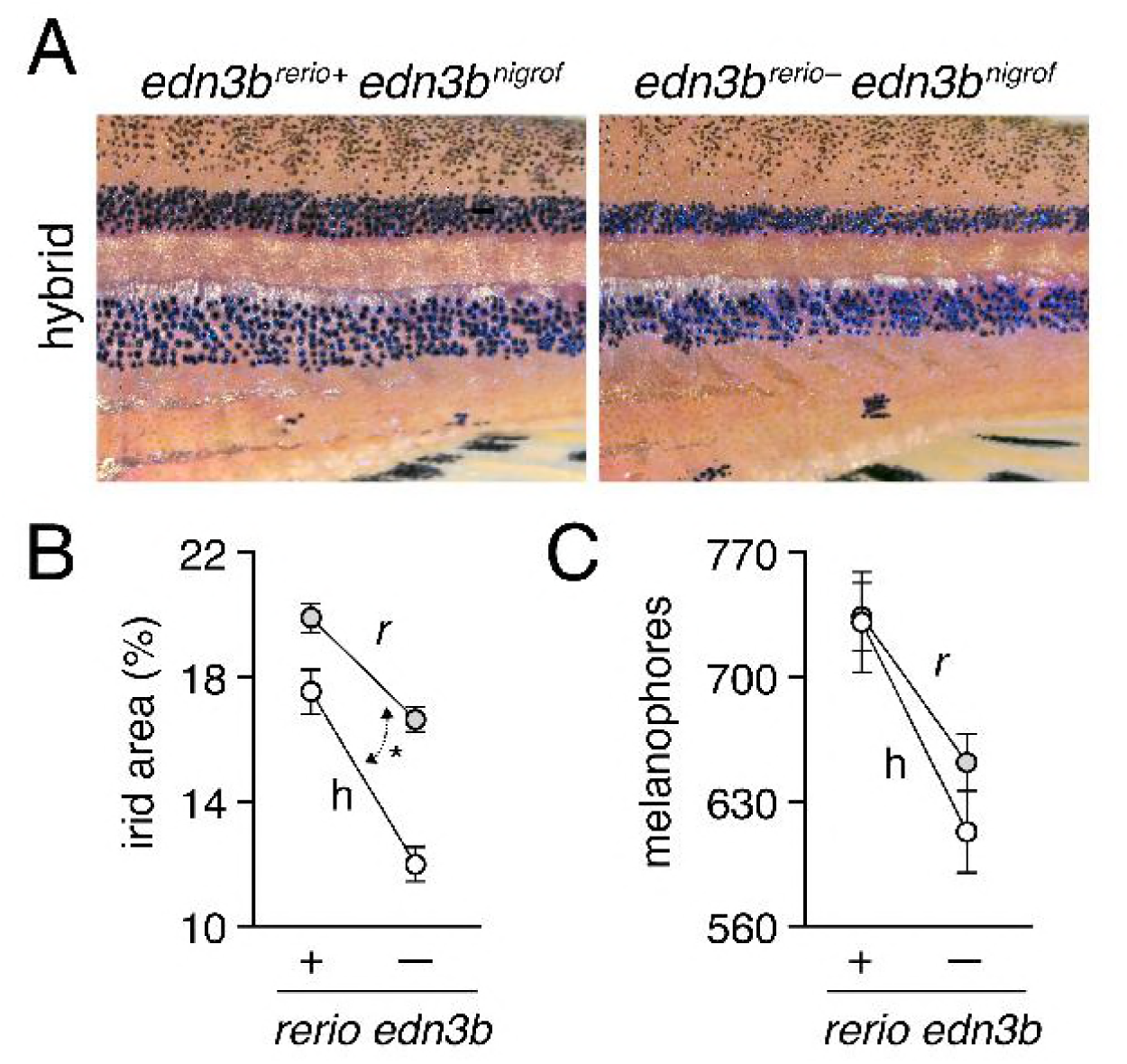
Hypomorphic edn3b allele in *D. nigrofasciatus* relative to *D. rerio*. (A) Interspecific hybrids carrying either *D. rerio* wild-type *edn3b* allele (left) or mutant *edn3b* allele (right). Carriers of the mutant allele tended to have narrower stripes and reduced iridophore coverage overall. (B) Hybrids (h) had reduced coverage by dense iridophores of interstripes (total percent of flank) compared to *D. rerio* (*r*) overall (*F*_1,59_=21.7, *P*<0.0001). Iridophore coverage was also reduced by substitution of a *D. rerio edn3b* mutant allele (−) for the *D. rerio* wild-type allele (+; *F*_1,59_=101.6, respectively; both *P*<0.0001), but this effect was more pronounced in hybrids, resulting in a significant background x allele interaction (*F*_1,59_=6.5, *P*<0.05; double headed arrow, different slopes). (C) Numbers of hypodermal melanophores were affected by background and *D. rerio* allele (*F*_1,61_=23.5, *F*_1,61_=24.7, respectively; both *P*<0.0001), but a background x allele interaction was non-significant (*F*_1,59_=1.0, *P*=0.3). Plots show least squares means±SE after controlling for significant effects of SL (*P*<0.05, *P*<0.0001, respectively). Sample sizes (*N*): 17 *D. rerio* (+); 24 *D. rerio* (−); 10 hybrids (+); 13 hybrids (−).

Two other genes, *augmentor*-*α1a* and *augmentor*-*α1b*, encoding secreted ligands for Leukocyte tyrosine kinase (Ltk), promote iridophore development in *D. rerio* and together have a mutant phenotype resembling *D. nigrofasciatus* [40,41]. Iridophore coverage in hybrids carrying *D. rerio* mutant alleles of *augmentor*-*α1a* and *augmentor*-*α1b* did not differ from siblings carrying *D. rerio* wild-type alleles (*F*_1,12_=0.01 *P*=0.9; *F*_1,11_,=0.3 *P*=0.6), highlighting specificity of the noncomplementation phenotype observed for *edn3b*.

### Reduced *edn3b* expression in skin of *D. nigrofasciatus* compared to *D. rerio* owing to *cis*-regulatory differences

A hypomorphic allele of *edn3b* in *D. nigrofasciatus* could result from changes in protein sequence conferring diminished activity, or changes in regulation causing reduced Edn3b abundance. The inferred protein sequence of *D. nigrofasciatus* Edn3b did not have obvious lesions (e.g., premature stop codon, deletions or insertions), and the 21 amino acid mature peptide was identical between species.

We therefore asked whether *D. nigrofasciatus edn3b* might be expressed differently than the *D. rerio* allele. Presumably owing to low overall levels of expression, *edn3b* transcripts were not detectable by *in situ* hybridization, and transgenic reporters utilizing presumptive regulatory regions amplified by PCR (~5 kb) or contained within bacterial artificial chromosomes (~190 kb containing ~105 kb upstream to the transcriptional start) failed to yield detectable fluorescence, precluding the assessment of spatial variation in gene expression. Nevertheless, quantitative RT-PCR on isolated skins of post-embryonic larvae indicated *edn3b* expression in *D. nigrofasciatus* at levels approximately one-quarter that of *D. rerio* (Figure 5A). Expression of *edn3b* was similarly reduced in the sister species of *D. nigrofasciatus, D. tinwini*, which has fewer melanophores and iridophores than *D. rerio*, and a spotted rather than striped pattern (Figure S4) [1,9].

**Figure 5.**
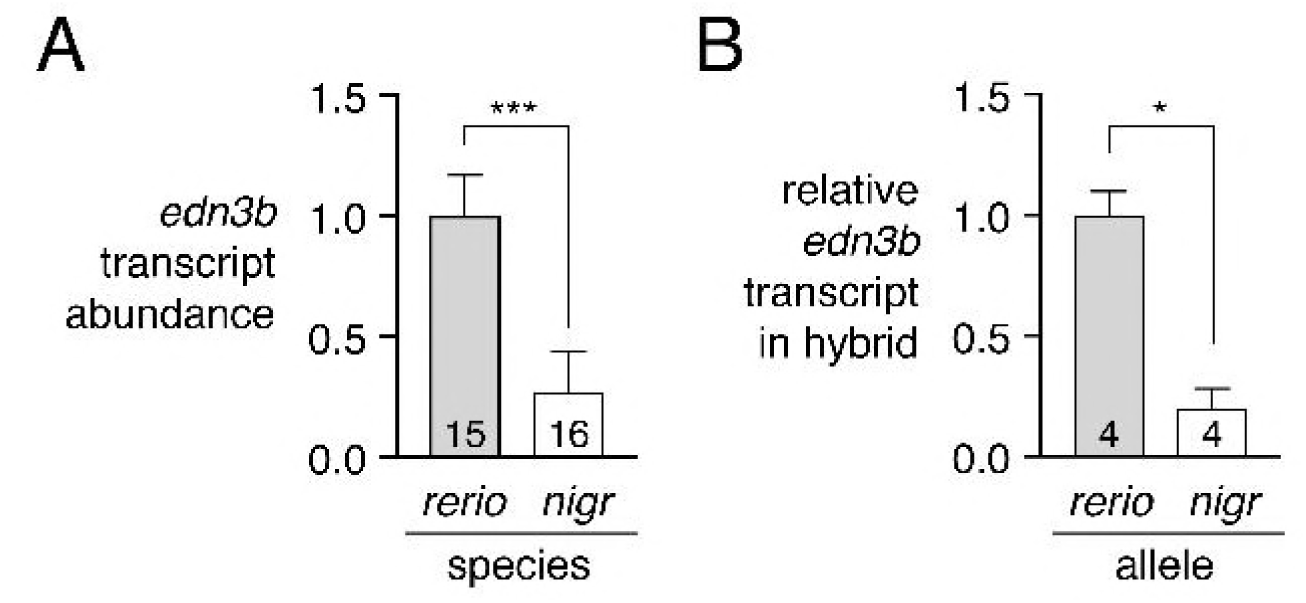
Lower expression of *D. nigrofasciatus edn3b* relative to *D. rerio edn3b*. (A) *edn3b* was expressed at lower levels in skin of *D. nigrofasciatus* than *D. rerio* (*F*_1,29_=48.6, *P*<0.0001) during adult pattern formation. (B) In hybrid fish, *D. nigrofasciatus edn3b* alleles were expressed at lower levels than *D. rerio edn3b* alleles (paired *t*_3_=4.6, *P*<0.05). Shown are means±SE. Values within bars indicate sample sizes.

This difference in *edn3b* expression raised the possibility that *cis*-regulatory factors (e.g, transcription factor binding sites, chromatin accessibility at *edn3b*) have been altered between *D. rerio* and *D. nigrofasciatus*. To test this idea, we compared expression of *D. rerio* and *D. nigrofasciatus edn3b* alleles in the common trans-regulatory background of *D. rerio* x *D. nigrofasciatus* hybrids. Allele-specific quantitative RT-PCR revealed approximately one-quarter the abundance of *D. nigrofasciatus edn3b* transcript compared to *D. rerio edn3b* transcript (Figure 5B). These observations suggest that species differences in *edn3b* result at least in part from *cis*-regulatory variation that drives lower levels of *edn3b* transcription in *D. nigrofasciatus* compared to *D. rerio*.

### Edn3b promotes increased iridophore coverage and secondarily affects melanophore pattern in *D. nigrofasciatus*

If lower expression of *edn3b* contributes to the difference in pigment pattern between *D. nigrofasciatus* and *D. rerio*, then expressing *edn3b* at higher levels in *D. nigrofasciatus* should generate a pattern converging on that of *D. rerio*. To test this prediction, we constructed stable transgenic lines in both species to express *D. rerio* Edn3b linked by viral 2A sequence to nuclear-localizing Venus, driven by the ubiquitously expressed heat-shock inducible promoter of *D. rerio hsp70l* [16,23]. We then reared *D. rerio* and *D. nigrofasciatus* transgenic for *hsp70l:edn3b*-*2a*-*nlsVenus*, and their non-transgenic siblings, under conditions of repeated heat shock during adult pigment pattern formation.

Heat-shock enhanced expression of Edn3b increased iridophore coverage in *D. nigrofasciatus* as compared to *D. rerio* or non-transgenic siblings of either species (Figure 6A,E). Excess Edn3b failed to increase total numbers of melanophores in *D. nigrofasciatus* (Figure 6B). Nevertheless melanophores were differentially distributed in these fish, as *D. nigrofasciatus* overexpressing Edn3b had about twice as many cells localizing in a secondary ventral stripe (2V), and a correspondingly reduced number of cells in the primary ventral stripe (1V), as compared to control siblings (Figure 6D). In *D. rerio*, total melanophore numbers were increased by Edn3b overexpression though melanophore distributions were not differentially affected between its normally complete stripes (Figure 6B,C,E).

**Figure 6.**
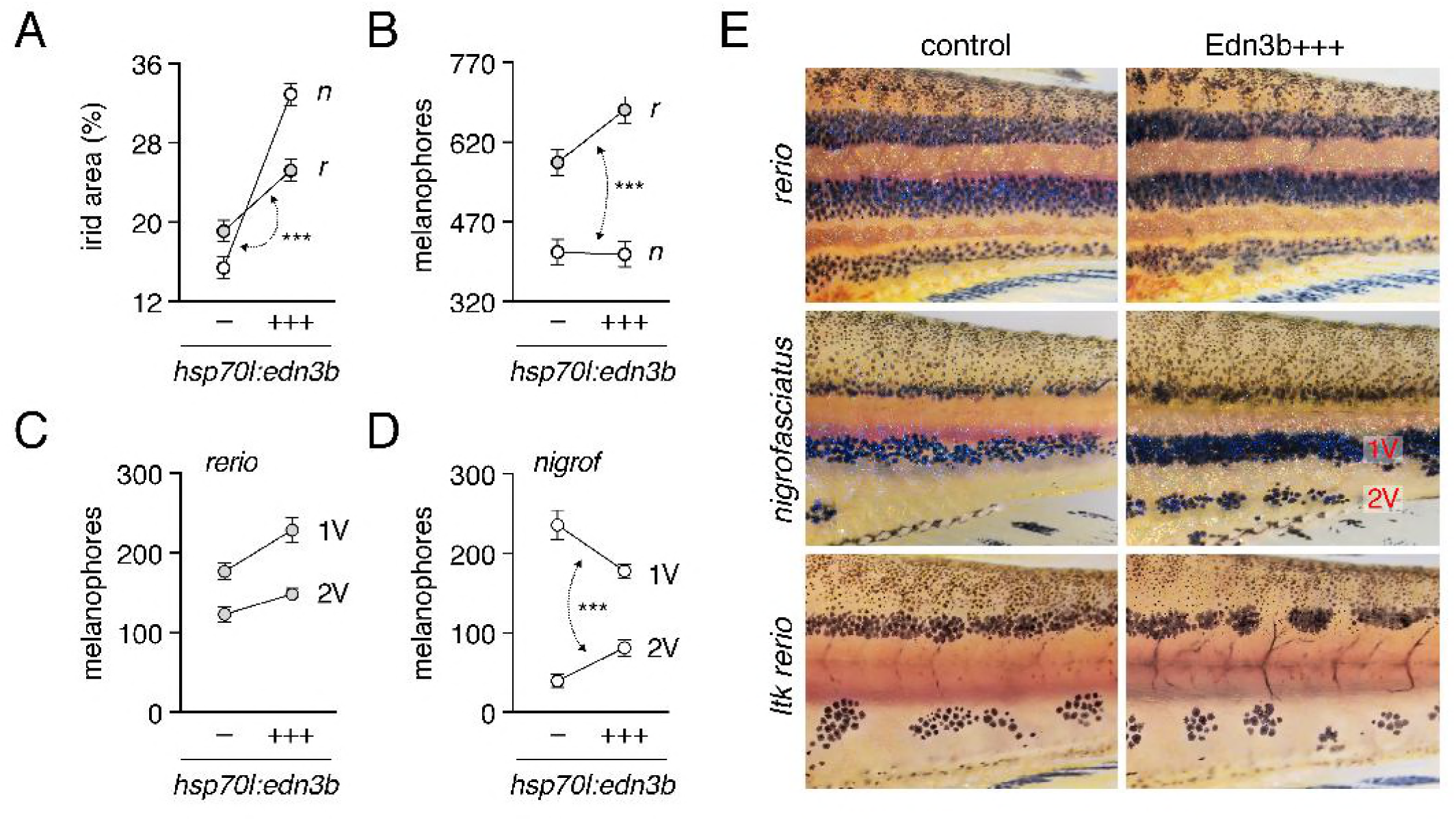
Edn3b increases iridophore coverage in both species and affects melanophore distribution indirectly in *D. nigrofasciatus*. (A) In both *D. rerio* (*r*) and *D. nigrofasciatus* (*n*), relative areas of the flank covered by interstripe (dense) iridophores was increased in response to Edn3b overexpression (+++) as compared to non-transgenic (−) sibling controls treated identically. The response to Edn3b overexpression was more pronounced in *D. nigrofasciatus* than in *D. rerio* (species x transgene interaction, *F*_1,55_=26.49, *P*<0.0001; double headed arrow, different slopes) (B) Edn3b overexpression increased total numbers of hypodermal melanophores in *D. rerio* but not *D. nigrofasciatus* (species x transgene interaction, *F*_1,55_=4.7, *P*<0.05). (C,D) Distributions of melanophores in ventral primary (1V) and ventral secondary (2V) stripes of *D. rerio* (C) and *D. nigrofasciatus*. In *D. nigrofasciatus*, Edn3b overexpression did not increase the total numbers of melanophores in these stripes (*F*_1,28_=0.4, *P*=0.5) but did result in a reallocation of melanophores from 1V to 2V (paired comparison within individuals, stripe position x transgene interaction (*F*_1,26_=71.0, *P*<0.0001). All plots shows means±SE. (E) Top and middle panels, Phenotypes of each species with and without Edn3b overexpression. Lower panels, Iridophore-free *Itk* mutant *D. rerio* in which Edn3b overexpression did not affect the numbers of melanophores (*F*_1,27_=1 .5, *P*=0.2) or their allocation between regions (paired comparison within individuals, stripe position x transgene interaction, *F*_1,27_=1.5, *P*=0.2). Sample sizes (*N*): 15 *D. rerio* (−); 15 *D. rerio* (+++); 15 *D. nigrofasciatus* (−); 15 *D. nigrofasciatus* (+++); 15 *Itk* mutant *D. rerio* (−); 15 *Itk* mutant *D. rerio* (+++).

The rearrangement of a constant number of melanophores in *hsp70l:edn3b*-*2a*-*nlsVenus D. nigrofasciatus*, and a requirement for interactions between iridophores and melanophores during normal stripe formation in *D. rerio* [15,16,23], raised the possibility that Edn3b effects on melanophores might be largely indirect, and mediated through iridophores. If so, we predicted that in a background entirely lacking iridophores, *hsp70l*:Edn3b should fail to affect melanophore numbers or distribution. We therefore generated fish transgenic for *hsp70l:edn3b*-*2a*-*nlsVenus* and homozygous for a mutant allele of *leucocyte tyrosine kinase* (*ltk*), which acts autonomously to promote iridophore development [15,40]. Consistent with iridophore-dependent Edn3b effects, neither melanophore numbers nor melanophore distributions differed between transgenic and non-transgenic siblings (Figure 6E, bottom panels).

These findings support a model in which lower expression of *edn3b* in *D. nigrofasciatus* results in diminished coverage by iridophores and a resulting failure of melanophores to more fully populate the secondary ventral stripe, as compared to *D. rerio*.

### Iridophore proliferation is curtailed in *D. nigrofasciatus* and *edn3b* mutant *D. rerio*

Finally, we sought to better understand the cellular bases for Edn3 effects on iridophore populations in *D. rerio* and *D. nigrofasciatus*. Given roles for Edn3 in promoting the proliferation of avian and mammalian neural crest cells and melanocytes [42–44], we hypothesized that *Danio* Edn3b normally promotes iridophore proliferation and we predicted that such proliferation would be curtailed in both *edn3b* mutant *D. rerio* and in *D. nigrofasciatus*.

To test these predictions, we examined iridophore behaviors by time-lapse imaging of larvae in which iridophores had been labeled mosaically with a *pnp4a:palm*-*mCherry* transgene. We detected iridophore proliferation in stripe regions, where these cells are relatively few and dispersed, and also within interstripes, where iridophores are densely packed (Figure 7). Proliferation of stripe-region iridophores was ~10-fold greater than that of interstripe iridophores. But within each region, iridophores of wild-type (*edn3b*/+) *D. rerio* were more likely to divide than were iridophores of *edn3b* mutants. Iridophores of *D. nigrofasciatus* had a proliferative phenotype intermediate to those of wild-type and *edn3b* mutant *D. rerio*. We did not observe gross differences in the survival or migration of iridophores across genetic backgrounds. These findings are consistent with Edn3b-dependent differences in iridophore proliferation affecting pattern formation both within *D. rerio*, and between *D. rerio* and *D. nigrofasciatus*.

**Figure 7.**
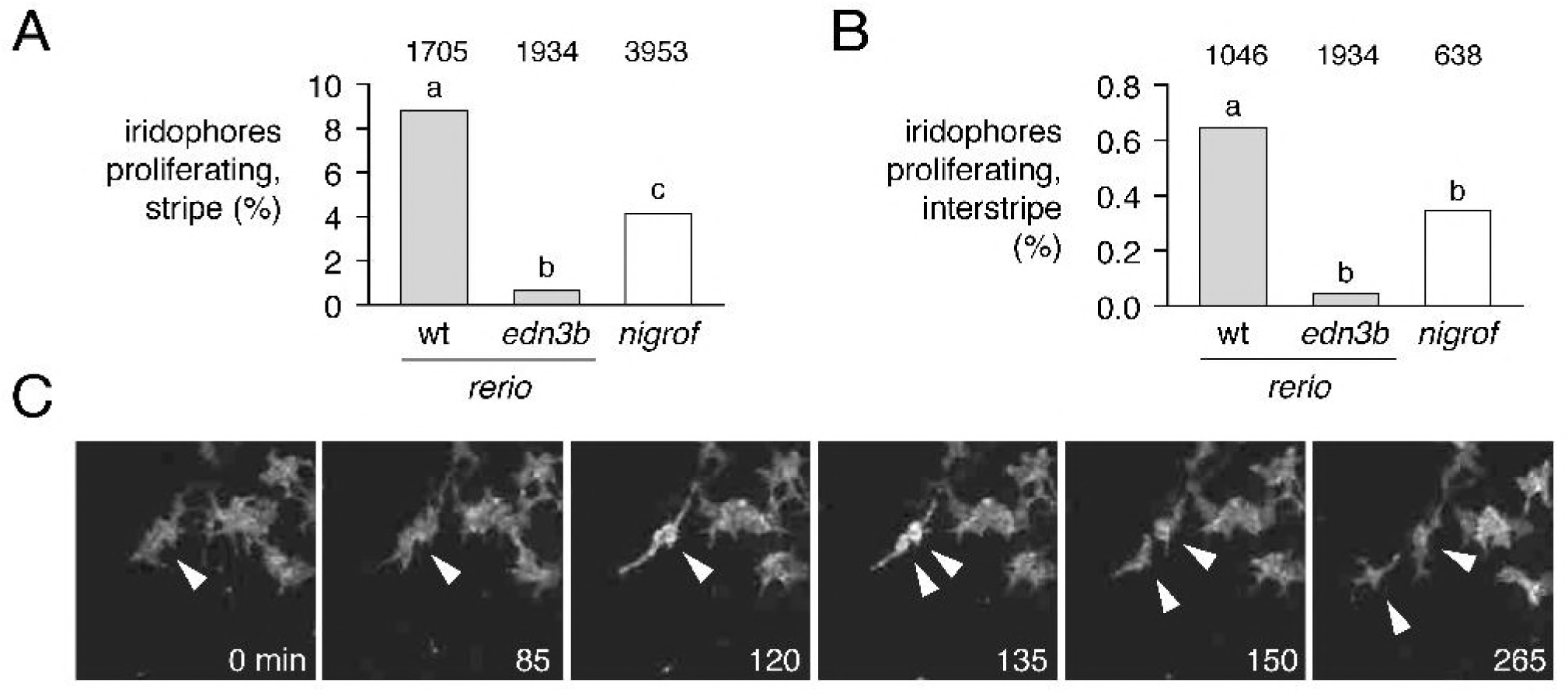
Reduced iridophore proliferation in edn3b mutant *D. rerio* and *D. nigrofasciatus*. (A) Among loosely organized iridophores of prospective stripe regions, the percent of individual cells dividing during time-lapse imaging (15 h total duration) was greatest in *edn3*/+ (wt) *D. rerio* and markedly reduced in sibling *edn3b* mutant *D. rerio* as well as *D. nigrofasciatus* (logistic regression: genotype, χ^2^=77.5, d.f.=2, *P*<0.0001; SL, χ^2^=77.6, d.f.=1, *P*<0.0001). (B) These same trends were evident for densely arranged iridophores of interstripes, though proliferation overall was reduced in comparison to stripe iridophores (genotype, χ^2^=13.7, d.f.=1, *P*<0.005; SL, χ^2^=31.9 d.f.=1 *P*<0.0001). Values above bars indicate total numbers of iridophores examined. Preliminary analysis did not reveal significant variation among individual larvae, the cells of which were pooled for final analyses (larval numbers: 8 *edn3*/+ *D. rerio*; 8 *edn3b* mutant *D. rerio*; 9 *D. nigrofasciatus*). Different letters above bars indicate genotypes that differed significantly from one another in pairwise comparisons of odds ratios (all *P*<0.005). (C) Stills from time-lapse video illustrating a single iridophore (arrowhead) within a prospective stripe region that partially rounds up by 120 min and then divides.

## Discussion

Towards a fuller understanding of pigment pattern diversification, we have analyzed cellular and genetic bases for differences in adult pattern between *D. rerio* and *D. nigrofasciatus*. Our study uncovers evolutionary changes in iridophore behavior between these species, identifies endothelin signaling as a candidate pathway contributing to these changes, and provides new insights into the evolution of endothelin genes and functions.

### Evolution of iridophore behaviors and impact on pattern reiteration

An important finding of our analyses is that evolutionary alterations in iridophore behavior can drive species differences in overall pattern. *D. rerio* and *D. nigrofasciatus* have relatively similar complements of iridophores during early stages of adult pattern formation, but the two species subsequently diverge from one another. In *D. rerio*, iridophore clone sizes expanded markedly as the fish grew and secondary and tertiary interstripes were added, whereas this expansion— and pattern element reiteration—were curtailed in *D. nigrofasciatus*. The difference in clonal expansion reflected, at least in part, differences in iridophore proliferation as revealed by time-lapse imaging.

Prior efforts documented the essential function of iridophores in promoting melanophore stripe reiteration [16,23]. Here, we showed that enhancing the iridophore complement of *D. nigrofasciatus* by Edn3b overexpression was sufficient to reallocate melanophores from a well-formed primary ventral stripe into an otherwise vestigial secondary ventral stripe, resulting in a pattern more like that of *D. rerio*. This effect was probably mediated by interactions between iridophores and melanophores, as melanophores did not respond to the same transgene in the *ltk* mutant of *D. rerio*, which lacks iridophores. An indirect role for endothelin signaling in promoting melanophore stripe development has likewise been inferred from cell transplantation between wild-type and *ednrb1a* mutant *D. rerio* [15], despite expression of *ednrb1a* by newly differentiating melanophores [34] and a responsiveness of *D. rerio* melanoma cells to Edn3b in the absence of iridophores [45].

Our observations suggest that an early cessation of iridophore clonal expansion in *D. nigrofasciatus* has led to an earlier offset of interactions between iridophores and melanophores, and an attenuation of the stripe pattern in *D. nigrofasciatus*. In heterochronic terms, the *D. nigrofasciatus* patterns could thus be described as pedomorphic relative to an inferred ancestral state, and arising by progenesis, relative to overall somatic development [46]. That a temporal change in the availability of interactions with iridophores has cascading effects on pattern is reminiscent of observations for xanthophores: precocious widespread xanthophore development, and resulting xanthophore–melanophore interactions, are associated with fewer stripes and more uniform pattern in *D. rerio* and *D. albolineatus* [23]. These outcomes highlight the diversity of patterns that can arise from a common set of cellular interactions in response to evolutionary modifications to the temporal or spatial pattern of pigment cell appearance.

### A role for endothelin signaling in *Danio* pattern evolution

The numerous pigment mutants of *D. rerio* might be expected to include genes that have contributed to evolutionary diversification within *Danio*, particularly when patterns of mutants and species resemble one another. We found that *edn3b* mutants of *D. rerio* have fewer iridophores and pattern elements than wild-type *D. rerio*, similar to the naturally occurring pattern of *D. nigrofasciatus*. This similarity of final phenotype was presaged by similarity of developmental phenotype, as both *edn3b* mutant *D. rerio* and *D. nigrofasciatus* had reduced iridophore proliferation relative to wild-type *D. rerio*.

Our study provides several lines of evidence to support a model in which alterations affecting Edn3b have contributed to the species difference in pigmentation. First, hybrids of *D. rerio* and *D. nigrofasciatus* carrying a mutant *D. rerio* allele of *edn3b* had a more severe iridophore deficiency than heterozygous *D. rerio* carrying the same mutant allele, suggesting that the *D. nigrofasciatus* wild-type allele is weaker than the *D. rerio* wild-type allele. Second, *edn3b* overexpression was sufficient to increase iridophore coverage, and (indirectly) alter melanophore distributions in *D. nigrofasciatus* to a state more similar to that of *D. rerio*. Third, we found reduced expression of *edn3b* in skin of *D. nigrofasciatus* compared to *D. rerio* during adult pigment pattern formation. Fourth, species differences in expression of *edn3b* alleles were re-capitulated even in a shared hybrid genetic background, pointing to evolutionary change in *cis*-regulation of this locus. Both *D. nigrofasciatus* and *D. tinwini* exhibited lower levels of *edn3b* expression compared to *D. rerio* so regulatory alteration(s) likely occurred prior to divergence of *D. nigrofasciatus* and *D. tinwini*, or within the lineage leading to *D. rerio* itself. *cis*-regulatory evolution affecting abundance of a secreted ligand that acts on pigment cells to affect pattern is similar to xanthogenic factor Csf1a of *Danio* [23], melanogenic Kit ligand of stickleback [47], and some aspects of anti-melanogenic Agouti in deer mice [48].

Our findings support a role for *edn3b* in *Danio* pattern evolution yet they also point to roles for additional factors. For example, overexpression of Edn3b in *D. nigrofasciatus* increased the coverage of iridophores and allowed for some rearrangements of melanophores, but failed to entirely recapitulate the pattern of *D. rerio*. Indeed, melanophore numbers were unchanged in transgenic *D. nigrofasciatus*, in contrast to the larger overall numbers of melanophore in wild-type *D. rerio* and the still larger number of melanophores induced indirectly by Edn3b overexpression in *D. rerio* (Figure 6B). Thus, pigment pattern differences between these species are clearly polygenic, and it seems likely that additional loci, of the endothelin pathway or other pathways, will be identified as contributing to attenuated stripes and interstripes of *D. nigrofasciatus* compared to *D. rerio*.

The endothelin pathway has been implicated in naturally arising strain differences previously. Besides the spontaneous mutant alleles of mouse Edn3 and Ednrb that allowed the pathway to be first characterized molecularly [49,50], endothelin pathway genes or differences in their expression have been associated with tabby coloration in domestic and wild cats [51], melanocyte deficiency in ducks [52], white and hyper-melanistic variants of chicken [53–55] and the white mutant axolotl [56]. It is tempting to speculate that mild alleles of endothelin pathway genes or alterations that affect their expression have relatively few pleiotropic effects, particularly in *Danio*, in which functions of Edn3 paralogues have become subdivided between distinct classes of iridophores. Pigmentary phenotypes associated with this pathway may be particularly accessible targets for natural or artificial selection.

### Evolution of endothelin genes and functions

Finally, our investigation of Edn3b bears on our understanding of how the endothelin pathway and its functions have evolved. Endothelins were discovered for their roles in vasoconstriction and have since been identified to have a variety of functions [29]. In the context of pigmentation, endothelins and their receptors have been most extensively studied in mammals and birds, in which they regulate proliferation, migration, differentiation and survival at various points within the neural crest–melanocyte lineage [30,31,33]. In teleosts, our results in *Danio* suggest that Edn3 acts primarily to promote iridophore development, with only indirect effects on melanophores. By contrast, the salamander *Ambystoma mexicanum* requires *edn3* for the development of melanophores, xanthophores and iridophores [56–58] and such effects are not plausibly mediated through iridophores, which develop long after the requirement by melanophores and xanthophores is first manifested.

In teleosts, an additional round of whole genome duplication has resulted in extra genes as compared to non-teleost vertebrates [59–61]. Though many duplicated genes have been lost, those having roles in pigmentation, including genes of the endothelin pathway have been differentially retained [28,29,62–64], presumably owing to the partitioning of ancestral functions and the acquisition of new functions. Our finding that *edn3a* and *edn3b* are required by complementary subsets of iridophores is consistent with subfunctionalization of an ancestral locus required by all iridophores.

Given requirements for Edn3 in other species—and our findings in *Danio* that *edn3a* and *edn3b* are required by iridophores, *edn3b* is required only indirectly by melanophores, and neither locus is required by xanthophores—we can propose a model for functional evolution in which: (i) an ancestral vertebrate Edn3 locus promoted the development of all three classes of pigment cells in ectotherms (a situation currently represented by *A. mexicanum*); (ii) loss of iridophores and xanthophores in mammals and birds obviated an Edn3 role in these cell lineages; (iii) Edn3 functional requirements became limited to iridophores in the lineage leading to teleost fishes and then were further partitioned between iridophore populations, at least in *Danio*. Further testing of this scenario will benefit from analyses of additional anamniotes, including gar, which diverged from the teleost lineage prior to the teleost genome duplication [61,65] and might be expected to have an Edn3 requirement similar to that of *A. mexicanum*.

## Materials and Methods

### Ethics statement

All animal research was conducted according to federal, state and institutional guidelines and in accordance protocols approved by Institutional Animal Care and Use Committees at University of Washington, University of Virginia and University of Oregon. Anesthesia and euthanasia used MS-222.

### Fish stocks and rearing conditions

Fish were reared under standard conditions (14L:10D at ~28 °C) and staging followed [14]. *Danio rerio* were inbred wild-type WT(ABb), a derivative of AB^∗^. CRISPR/Cas9 mutants were induced in WT(ABb) (*edn3b^vp.r30c1^*) or ABC x TU (*edn3a^b1282^*, *edn3b^b1283^*). *Danio nigrofasciatus* was field-collected in Myanmar in 1998 [37] and maintained in the laboratory since that time. *Danio tinwini* was obtained from the pet trade in 2014. Transgenic lines *hsp70l:edn3b*-*2a*-*nlsVenus^vp.rt30^* and *hsp70l:edn3b*-*2a*-*nlsVenus^vp.nt2^* were generated in WT(ABb) and *D. nigrofasciatus* backgrounds, respectively. *augmentor*-*α1a*/+ and *augmentor*-*α1b*/+ *D. rerio* [41] were generously provided by E. Mo and S. Nicoli (Yale School of Medicine). *ltkj^9s1^* (*primrose*) is a spontaneous allele of *ltk* identified by S. Johnson, into which *hsp70l:edn3b*-*2a*-*nlsVenus^vp.rt30^* was crossed.

Fish were fed marine rotifers, brine-shrimp and flake food. Fish were allowed to spawn naturally or gametes were stripped manually for *in vitro* fertilization. Interspecific hybrids were generated by *in vitro* fertilization in both directions using *D. rerio* heterozygous for wild-type and *edn3b^vp.r30c1^* allele; progeny were reared through formation of juveniles patterns and then genotyped using primers to amplify *D. rerio* alleles by PCR from fin clips, followed by Sanger sequencing to identify carriers or WT(ABb) or *edn3b^vp.r30c1^* alleles. For *hsp70l*-inducible Edn3b transgenes, transgenic siblings and non-transgenic controls were reared from stages DR through J under conditions of repeated daily heat shock (38°C, 1 h) [16,23].

### CRISPR/Cas9 mutagenesis, transgenesis and clonal analyses

For CRISPR/Cas9 mutagenesis, 1-cell stage embryos were injected with T7 guide RNAs and Cas9 protein (PNA Bio) using standard procedures [66]. Guides were tested for mutagenicity by Sanger sequencing and injected fish were reared through adult stages at which time they were intercrossed to generate heteroallelic F1s from which single allele strains were recovered. CRISPR gRNA targets (excluding proto-spacer adjacent motif) were: *edn3a^b1282^*, GCCAGCTCCTGAAACCCCAC; *edn3bvp*^vp.r30c1^, GAGGATAAATGTACTCACTG; *edn3b^b1283^*, GGATAAATGTACTCACTGTG

For transgenesis, constructs were generated using the Tol2Kit and Gateway cloning [67] and injected by standard methods with *Tol2* transposase mRNA [68]. For Edn3b-containing transgenes, F0 mosaic adults were screened for germline transmission and progeny tested for *hsp70l*-induction of linked fluorophore. Clonal analyses used mosaic F0 larvae and limiting amounts of *pnp4a:palmEGFP* transgene to insure that integrations were rare between and within individuals so that only single clones were likely to be labeled [35,36]. Sparsity of transgene+ embryos and similarity of starting clone sizes within such embryos between species suggests that labeling was indeed clonal. Transgene+ individuals were imaged at stages PR_+_ and J_++_.

### Quantitative RT-PCR

For assessing *edn3b* transcript abundance across species, skins were harvested from stage-matched *D. rerio*, *D. nigrofasciatus* and *D. tinwini* and total RNAs isolated by Trizol (ThermoFisher) extraction as previously described [23]. First strand cDNAs were synthesized with iScript and oligo-dT priming (BioRad) and analyzed on an ABI StepOne Plus real time PCR instrument using custom designed Taqman probes against target sequence shared by *D. rerio* and *D. nigrofasciatus* (identical to *D. tinwini*). *edn3b* expression was normalized to that of *rpl13a*; normalization to a conserved *actb1* amplicon (ThermoFisher assay ID #Dr03432610_m1) yielded equivalent results in pilot analyses (not shown). Expression levels were assessed using the 2^−ΔΔCt^ method [69] with *D. rerio* expression levels set to 1. Comparisons of species differences in expression were repeated 4 times (with 2–4 biological replicates each) using matched stages of fish between DR_+_ and J. We did not detect significant differences between replicates/stages, or species x replicate/stage interactions, and so present normalized values across all replicates in the text. For analyzing allele-specific expression in hybrids, custom Taqman probes were designed to amplify an *edn3b* target from both species alleles, or from only *D. rerio* (*Dr*) or *D. nigrofasciatus* (*Dn*). Amplifications of *Dr* and *Dn* probes were normalized to that of the *Dr*, *Dn* probe. Hybrid samples included a total of 4 biological replicates. Primers (F, R) and target probes (T) were: *edn3b* (AIWR3Z6): F-CAGAGAAT GT GTTT ATT ACT GT CATTTGGG, R-CCAAGGT GAACGT CCT CT CA, P-FAM-CTGGGATCAACACCCCACAACG; *edn3b*(AI20TXP, *Dr*): F-TGGTGGTTCCAGCAGTGTTG, R-TGTGAGCGTGTGATGCTGAA, P-FAM-CAAGCTTCGCTTCTTTC; *edn3b*(AI1RVRH, *Dn*): F-GCTCTTTTGCTAATTGTGAGTTTGGT, R-ACCAGAGAAGACTGGAGATGAGT, P-FAM-CT CCTGCACTT GAAAAC; *rpl13a* (*Dr*, *Dn*): F-CAGAGAAT GT GTTT ATT ACT GT CATTTGGG, R-CCAAGGTGAACGTCCTCTCA, P-FAM-CTGGGATCAACACCCCACAACG. Accession for *D. nigrofasciatus edn3b* is pending [submission #2127550].

### Imaging

Images were acquired on: Zeiss AxioObserver inverted microscopes equipped either with Axiocam HR or Axiocam 506 color cameras or a Yokogawa laser spinning disk with Evolve camera, and an AxioZoom v16 stereomicroscope with Axiocam 506 color camera, all running ZEN blue software. An Olympus SZX12 stereomicroscope with Axiocam HRc camera and Axiovision software was additionally used for some imaging. Images were corrected for color balance and adjusted for display levels as necessary with all treatments or species within analyses treated identically. Images of swimming fish were captured with a Nikon D800 digital SLR equipped with Nikon AF-S VR Micro-Nikkor f2.8 IF/ED lens.

Counts of melanophores and coverage by iridophores used regions of interest defined dorsally and ventrally by the margins of the flank, anteriorly by the anterior insertion of the dorsal fin and posteriorly by the posterior insertion of the anal fin. Only hypodermal melanophores contributing to stripes were included in analyses; dorsal melanophores and melanophores on scales were not considered. For assessing iridophore coverage, total areas covered by dense interstripe iridophores were estimated as these account for the majority of total hypodermal iridophores and areas covered by sparse iridophores within stripe regions could not be reliably estimated from brightfield images. Cell counts and area determinations were made using ImageJ. Time-lapse analyses of iridophore behaviors followed [21] and were performed for 15 h with 5 min frame intervals on *D. nigrofasciatus* as well as *D. rerio* siblings homozygous or heterozygous for *edn3b^vp.r30c1^*. Individual genotypes of larvae used for time-lapse imaging were assessed by Sanger sequencing across the induced lesion.

### Statistical analysis

All statistical analyses were performed using JMP 14.0.0 statistical analysis software (SAS Institute, Cary NC) for Apple Macintosh. For linear models residuals were examined for normality and homoscedasticity and variables transformed as necessary to meet model assumptions [70].

## Acknowledgements

Supported by NIH R35 GM122471 to D.M.P. and NIH P01 HD22486 and NIH R25 HD070817 to J.S.E. Thanks to A. Schwindling and other Parichy lab members for technical assistance with experiments and fish rearing.

---

**Figure S1.**
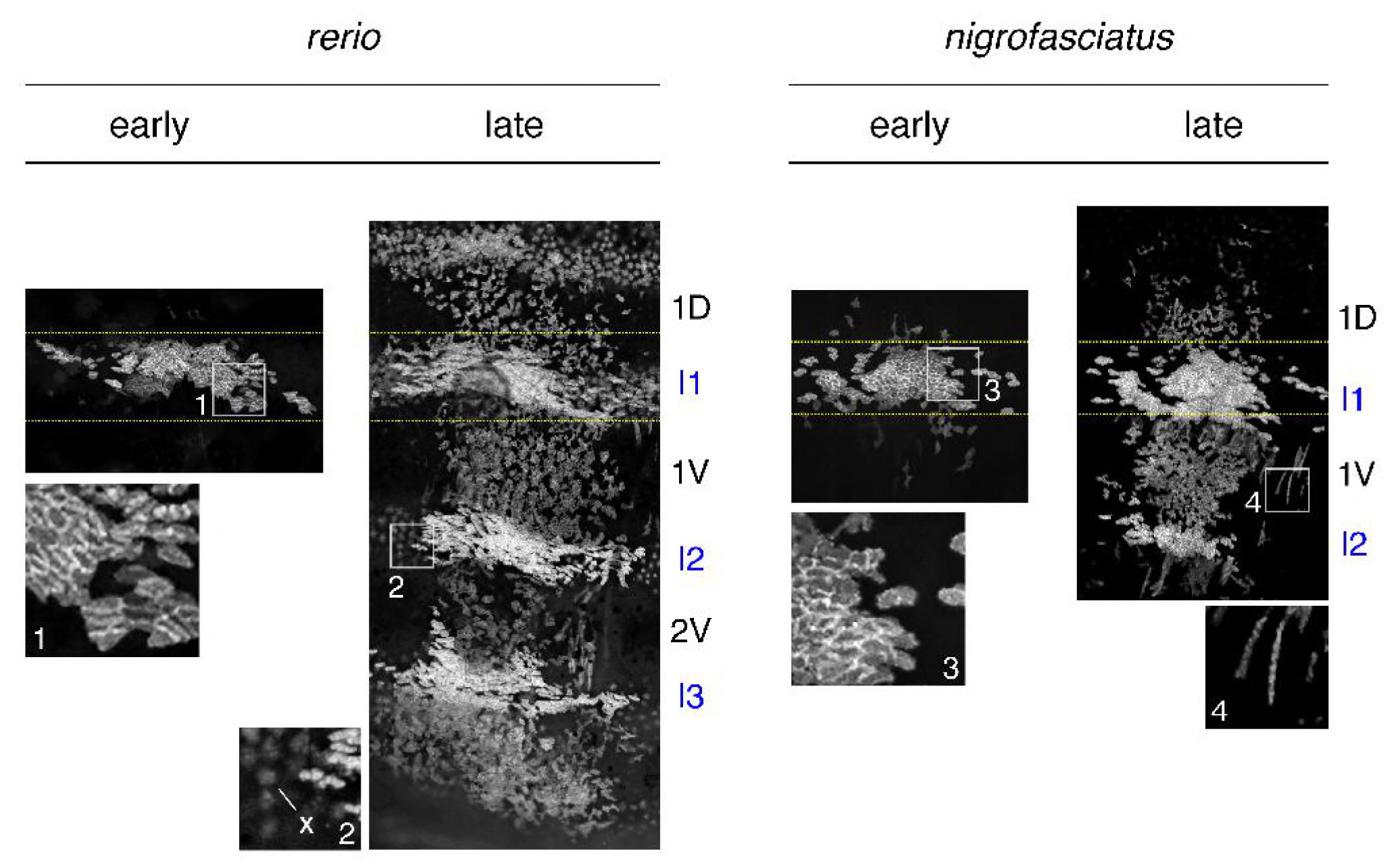
Expansion of iridophore clones differs between *D. rerio* and *D. nigrofasciatus*. Representative images for individuals of each species mosaic for iridophore reporter *pnp4a:palmEGFP* at an early stage of pattern formation, and a late stage, once patterns were complete. Dashed yellow lines indicate approximate regions of correspondence between early and late images and I1–I3 indicate primary through tertiary interstripes, if present; 1D, 1V, 2V indicate positions of stripes, if present. In each species, iridophores were present within interstripes, where they were densely packed, and within stripe, where they were loosely arranged. Inset 1, clonal derived early iridophores in primary interstripe of *D. rerio*. Inset 2, In some individuals, autofluorescent xanthophores (x) were apparent but were distinguishable from iridophores by differences in shape. Inset 3, early iridophores of *D. nigrofasciatus*. Inset 4, Examples of spindle-shaped “type-L” iridophores [71] present at low abundance in each species.

**Figure S2.**
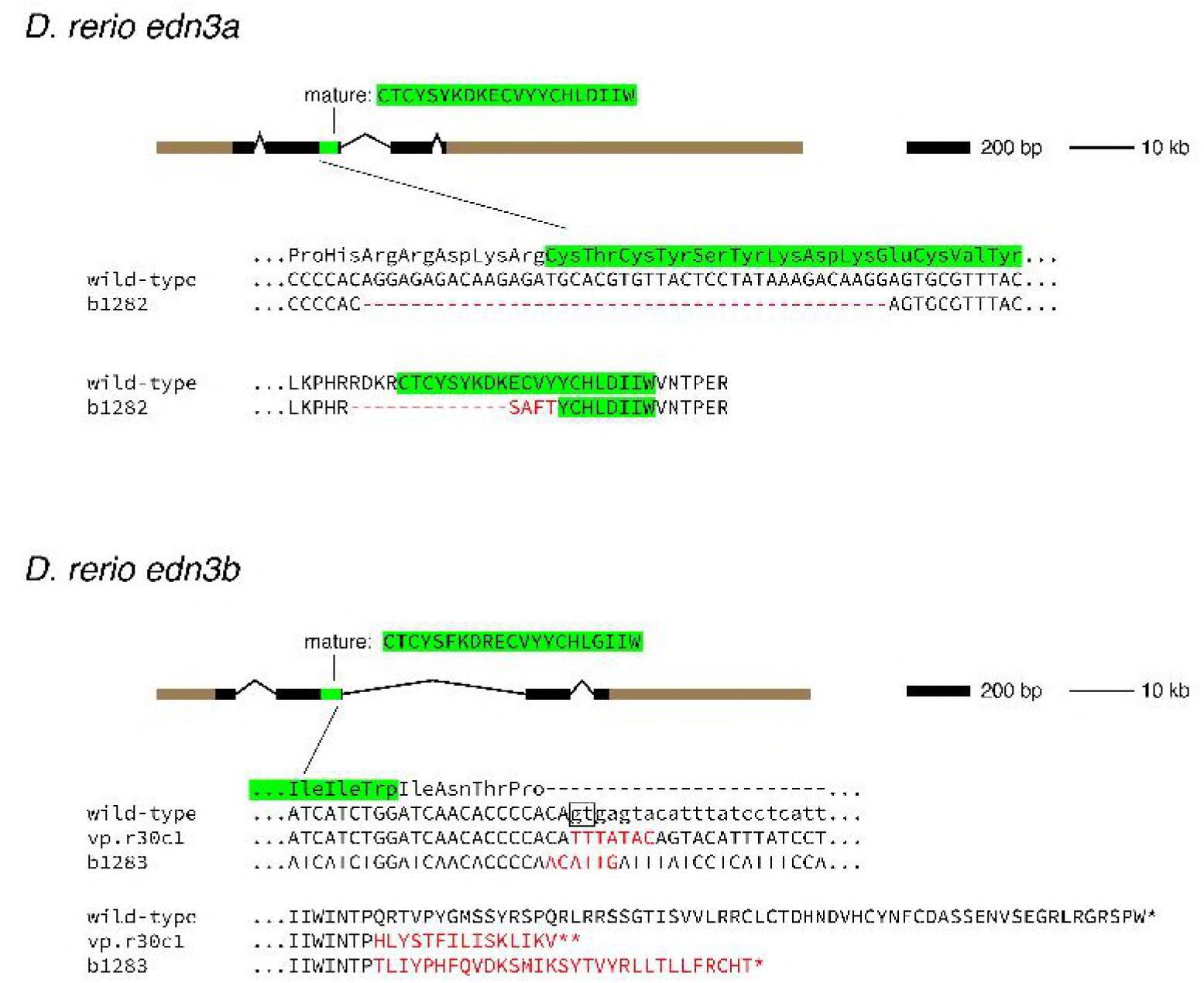
Induced mutations in *D. rerio* Edn3 loci. Panels show genomic structures of Edn3 loci with locations encoding the mature peptides (green) as well as local nucleotide and amino acid sequences. Untranslated regions are shown in brown. For *edn3a*, the *b1282* allele has a 43 bp deletion that removes 13 of 20 amino acids comprising the active Edn3a peptide, with the addition of 4 novel amino acids (red). For *edn3b*, two alleles were generated with deletions of existing nucleotides and insertion of new nucleotides (red) covering the splice donor site downstream of exon 2 (boxed), resulting in the addition of novel amino acids and premature stop codons (^∗^). Both *vp.r30c1* and *b1283* are likely to be loss-of-function mutations as their phenotypes were indistinguishable and also resembled independently derived *edn3b* alleles having similar lesions at the same target site [45]. Open reading frames are in upper case and intronic sequence in lower case.

**Figure S3.**
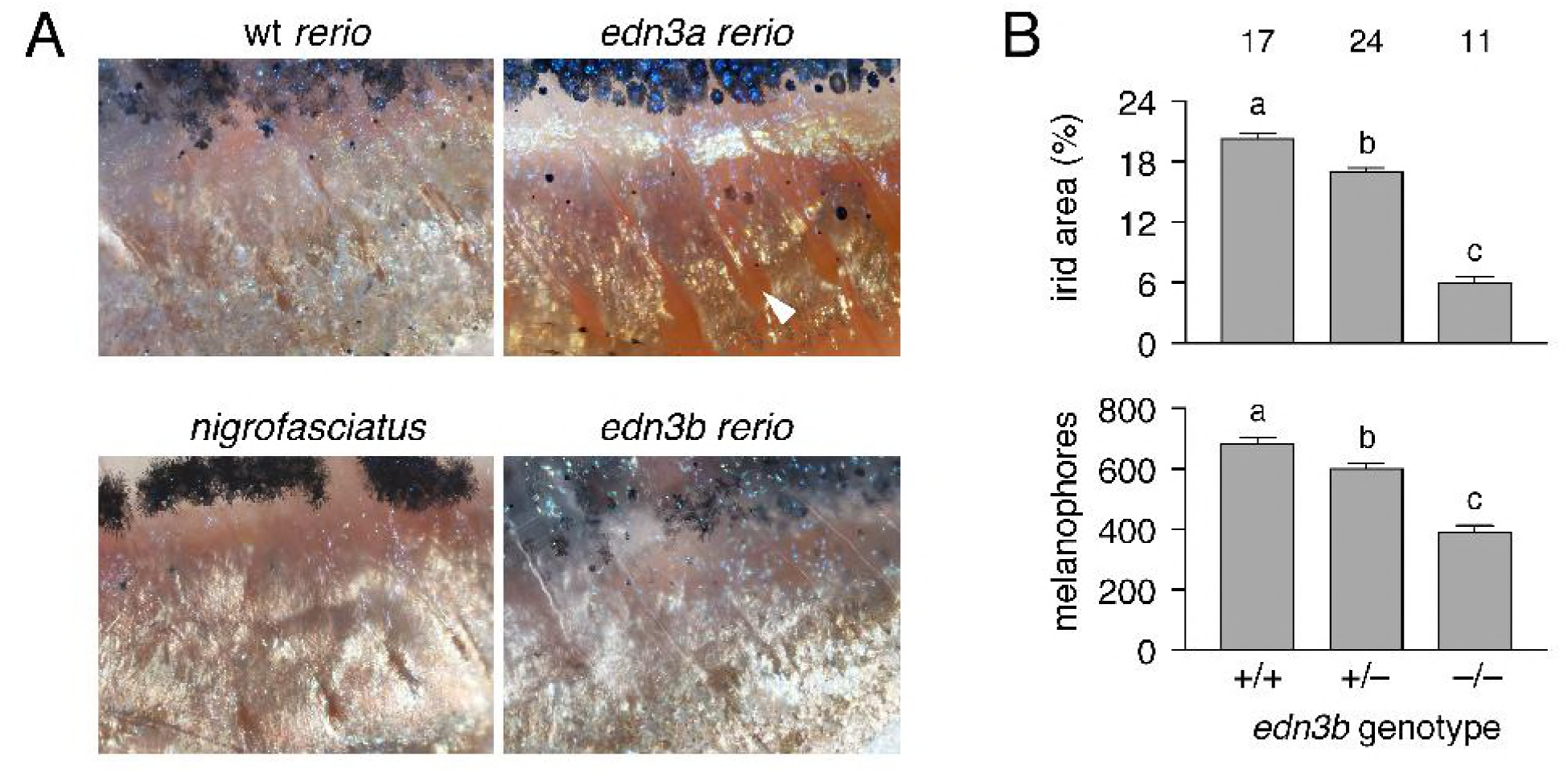
Pigment pattern defects of *edn3b* mutants but not *edn3a* mutants resemble *D. nigrofasciatus*. (A) Details of ventral patterns illustrating deficiency in peritoneal iridophores (arrowhead) in *D. rerio edn3a* mutants but not *edn3b* mutants or *D. nigrofasciatus*. (B) Defects in areas covered by iridophores and numbers of melanophores in heterozygous and homozygous *edn3b* mutant *D. rerio* (*F*_2,48_=292.6, *F*_2,48_=69.8, respectively; both *P*<0.0001). Shown are least squares means±SE after controlling for variation in standard length (SL; both *P*<0.0001). Different letters above bars indicate means significantly different in Turkey-Kramer post hoc comparisons. Values above bars indicate samples sizes.

**Figure S4.**
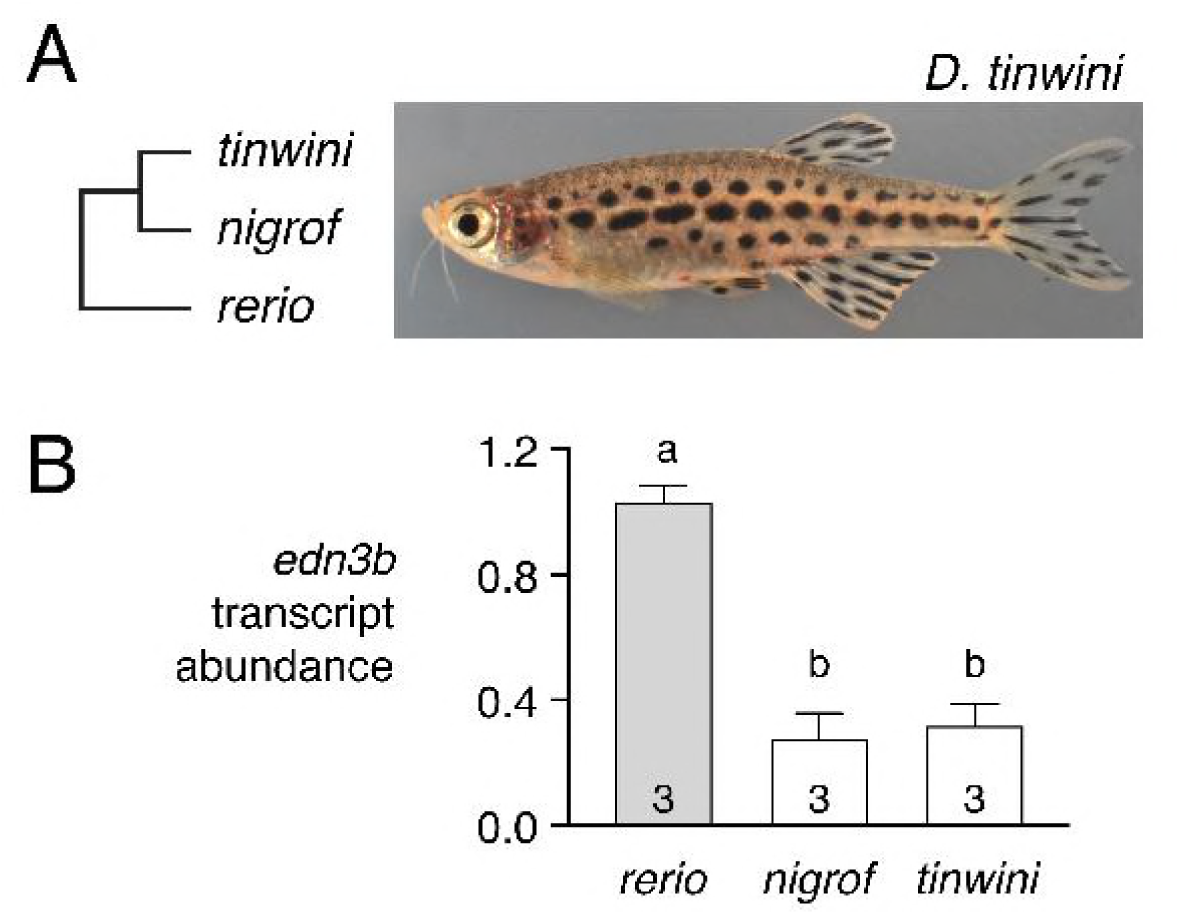
Reduced *edn3b* expression in *D. tinwini*compared to *D. rerio*. (A) Phylogenetic relationships [9] and pattern of *D. tinwini*. (B) Species differences in skin *edn3b* expression during adult pattern development (*F*_2,7_=48.2, *P*<0.0001). Shared letters indicate bars not significantly different in post hoc Turkey HSD comparisons of means (*P*>0.05). Numbers in bars indicate biological replicates.

